# The Semantics of Adjective Noun Phrases in the Human Brain

**DOI:** 10.1101/089615

**Authors:** Alona Fyshe, Gustavo Sudre, Leila Wehbe, Nicole Rafidi, Tom M. Mitchell

## Abstract

As a person reads, the brain performs complex operations to create higher order semantic representations from individual words. While these steps are effortless for competent readers, we are only beginning to understand how the brain performs these actions. Here, we explore semantic composition using magnetoencephalography (MEG) recordings of people reading adjective-noun phrases presented one word at a time. We track the neural representation of semantic information over time, through different brain regions. Our results reveal two novel findings: 1) a neural representation of the adjective is present during noun presentation, but this neural representation is different from that observed during adjective presentation 2) the neural representation of adjective semantics observed during adjective reading is reactivated after phrase reading, with remarkable consistency. We also note that while the semantic representation of the adjective during the reading of the adjective is very distributed, the later representations are concentrated largely to temporal and frontal areas previously associated with composition. Taken together, these results paint a picture of information flow in the brain as phrases are read and understood.

## 1. Introduction

Semantic composition, the process of combining small linguistic units to build more complex meaning, is fundamental to language comprehension. It is a skill that children acquire with amazing speed, and that adults perform with little effort. Still, we are just beginning to understand the neural processes involved in semantic composition, and the neural representation of composed meaning. Multiple studies have identified cortical regions exhibiting increased activity during increased semantic composition load ([1, 2, 3, 4, 5], to name only a few). Here we considered a different question: Where and when are semantic representations stored in preparation for, and used during semantic composition?

Semantic composition in the brain has been studied using semantically anomalous sentences [1, 2, 6], as well as in simple phrases [4, 5], typically by comparing the *magnitude* of brain activity between conditions (e.g. composing words into phrases vs. reading word lists). Several such studies have implicated right and left anterior temporal lobes (RATL and LATL) as well as ventro-medial prefrontal cortex (vmPFC) and left inferior frontal gyrus (IFG) in compositional processing [4, 7, 3]. Magnetoencephalography (MEG) studies have shown effects in these areas as early as 180ms post stimulus onset, until around 480 ms post stimulus onset [4, 7], which aligns well with the N400 effect observed in electroencephalography for semantically incongruent sentences [1]. Semantic composition effects have also been seen around 600ms post stimulus onset (P600)[8]. Though the P600 is more commonly associated with *syntactic* violations, it can be evoked by syntactically sound stimuli that violate a semantic constraint [2].

The computational (rather than neuroscientific) study of language semantics has been greatly influenced by the idea that word meaning can be inferred by the context surrounding a given word, averaged over many examples of the word’s usage [9, 10, 11, 12, 13]. For example, we might see the word *ball* with verbs like *kick, throw, catch*, with adjectives like *bouncy* or with nouns like *save* and *goal*. Context cues suggest meaning, so we can use large collections of text to compile statistics about word usage (e.g. the frequency of pairs of words), to build models of word meaning. These statistics are typically compressed using dimensionality reduction algorithms like singular value decomposition (SVD), as in latent semantic analysis (LSA) [14]. LSA and similar models represent each word with a vector. Together, the vectors of many words form a Vector Space Model (VSM).

The brain’s semantic representations can be studied by quantifying the information present in neural activity at particular cortical locations and times. For example, one can predict the word a person is reading based on their neural activity, by training a machine learning algorithm to predict the associated word vector [15, 16]. Such algorithms do not require large differences in brain activity between conditions, but rather leverage differences in the *spatio-temporal patterns* of neural activity, which may involve differences in signal in both the positive and negative direction in different areas of the brain at different times. Machine learning techniques have been used in a variety of settings to predict words from brain activity [15, 16, 17, 18, 19].

To our knowledge, the study presented here represents the first effort to study *semantic representations* of adjective-noun phrases using the fine time resolution offered by Magnetoencephalography (MEG). To study adjective-noun phrases in the brain, we traced the flow of information through time and brain space. The ability of these algorithms to predict the stimuli from MEG recordings is indicative of the information present in the underlying brain activity, and thus is indicative of the brain’s neural representation of that stimulus.

## 2. Materials and Methods

To examine the neural representations of adjective and noun semantics, we presented phrases consisting of an (adjective or determiner) followed by a noun. To maximize our probability of detecting the individual words of the phrase, we chose 8 nouns that can be easily predicted from MEG recordings [16]. We chose six adjectives to modulate the most predictable semantic qualities of the words (e.g. edibility, manipulability, size) [16]. We also paired nouns with “the” to isolate noun meaning. In total, there are 30 word pairs (phrases). For a full list, see the Appendix. Though some phrases start with a determiner (the), for simplicity, throughout this paper we will refer to all phrases as adjective-noun phrases. MEG data was recorded for 9 subjects (4 female), all neurologically healthy, right-handed, native English speakers with normal or corrected to normal vision.

Phrases are shown in rapid serial visual presentation (RSVP) format (See Figure 1). During each trial, the first word of the phrase appears on the display at 0 seconds, and is removed from the display at 500 ms. The noun appears at 800 ms and is removed at 1300 ms. To ensure subjects were engaged during the experiment, 10% of the stimuli were adjective-adjective pairs (oddballs), for which the subjects were instructed to press a button with their left hand. Neither the adjective-adjective trials, nor the adjective-noun trial immediately following the oddball were used for analysis. Excluding these omitted trials, each phrase was presented 20 times, and analysis was performed on the mean MEG time series over all 20 trials. The experiment was carried out in 7 blocks of approximately equal length, with the opportunity to rest between blocks.

**Figure 1:**
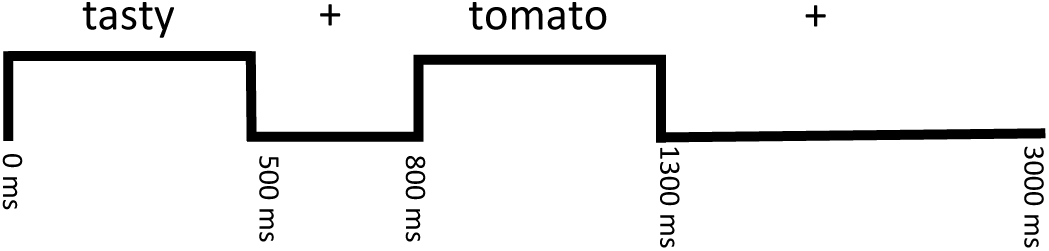
Stimulus presentation protocol. Adjective-noun phrases were presented one word at a time with a fixation cross both between words and between phrases. The first word of the phrase appears at 0 ms, and disappears at 500ms, the second word is visible 800 ms – 1300 ms. The first word of successive phrases are 3000 ms apart.

Because of the experimental setup, there is a strong correlation between the adjectives and nouns in our data. For example, the word “rotten” only appears with food words “tomato” and “carrot”. For this reason, we were careful to avoid analyses in which we seek to predict a property of the adjective, but might instead predict a correlated property of the noun (and vice versa). This is explained further in Section 2.6.

### 2.1. Data Acquisition and Preprocessing

All 9 subjects gave their written informed consent approved by the University of Pittsburgh (protocol PRO09030355) and Carnegie Mellon (protocol HS09-343) Institutional Review Boards. MEG data were recorded using an Elekta Neuromag device (Elekta Oy). As much as possible, we adhered to the best practices of MEG data collection[20]. The data were acquired at 1 kHz, high-pass filtered at 0.1 Hz and low-pass filtered at 330 Hz. Eye movements (horizontal and vertical eye movements as well as blinks) were monitored by recording the differential activity of muscles above, below, and beside the eyes. At the beginning of each session, the position of the subject’s head was recorded with four head position indicator (HPI) coils placed on the subject’s scalp. The HPI coils, along with three cardinal points (nasion, left and right pre-auricular), were digitized into the system to allow for head translation to register data collected in different blocks.

The data was preprocessed using the temporal extension of SSS (tSSS) [21] to remove artifacts and noise unrelated to brain activity. In addition, we used tSSS to realign the head position measured at the beginning of each block to a common location. The MEG signal was then low-pass filtered to 50 Hz to remove the contributions of line noise and down-sampled to 200 Hz. The Signal Space Projection method (SSP) [22] was used to remove signal contamination by eye blinks or movements, as well as MEG sensor malfunctions or other artifacts. The MEG data was parsed into trials, one for each phrase presentation. Each trial begins at the onset of the first word of the phrase, and ends 3000 ms later, for a total of 600 time points of data per sample (See Figure 1). MEG sensor amplitudes are known to drift with time, so we corrected each trial by subtracting from every sensor the mean signal amplitude during the 200ms before stimulus onset. During behavioral tests it was found that phrases containing the noun “thing” were inconsistently judged by human subjects, and so the 8 phrases containing the noun “thing” were omitted from further analysis, leaving a total of 30 phrases for analysis.

After processing, the MEG data for each subject consisted of 20 repetitions for each of the 30 phrases. Each repetition has a 600 dimensional time series for each of the 306 sensors. For each subject, we averaged all 20 repetitions of a given phrase to create one data instance per phrase, 30 instances in all. The dimensions of the final data matrix for each subject were 30 *×* 306 *×* 600.

### 2.2. Source Localization

In order to transform MEG sensor recordings into estimates of neural activity localized to areas of the brain, we used a multi-step process. First, Freesurfer (http://surfer.nmr.mgh.harvard.edu) was used to construct a 3D model of each subject’s brain, based on a structural MRI. Freesurfer was used to segment the brain into ROIs based on the Desikan-Killiany Atlas. Then the Minimum Norm Estimate method [23] was used to generate estimates of sources on the cortical sheet, spaced 5mm apart. The noise covariance matrix was estimated using approximately 2 minutes of MEG recordings collected without a subject in the room (empty room recordings) either directly before or after the subject’s session. Source localization resulted in approximately 12000 sources per subject derived from the 306 MEG sensor signals.

### 2.3. Prediction Tasks

We used two prediction tasks to track the words representations while reading adjective-noun phrases. In each case, the task is to predict the stimulus from the MEG recording. Differences in the time course of prediction accuracy for each task allows us to probe the information available during adjective-noun phrase comprehension. The tasks are:

1. **Predicting Adjective Semantics:** Predict the identity of the first word in the phrase (any of the 6 adjectives or the word “the”)
2. **Predicting Noun Semantics:** Predict the identity of the noun (one of 8).

For both tasks, we trained models to predict the dimensions of a VSM vector from MEG recordings. We then predict word identity based on the similarity of the predicted vectors to the corresponding true VSM vectors. A more detailed description follows in Section 2.4. For each of the words in our study, we used word vectors that are based on the sentence dependency relationships for each word of interest, averaged over a very large number of sentences [24]. For more details on the vectors see the Appendix. We use the first *m* = 100 SVD dimensions to summarize the dependency statistics, which represents a reasonable trade-off between computation time and accuracy [24].

### 2.4. Prediction Framework

To study adjective-noun composition in the brain we devised a simple prediction framework. Cross-validation was performed independently for each subject, wherein 2 of the 30 phrases are withheld during training, and subsequently used to test the framework’s predictions. This hold out and test procedure was repeated multiple times (see Section 2.6). For the prediction tasks described in Section 2.3, every stimulus phrase is represented by two vectors from a VSM, which represent the semantics of either the adjective or noun. The elements of this vector are the targets in the prediction framework (*s*^(*k*)^ in Equation 1).

We created a data matrix *X ∈* ℝ^*N×P*^ where *N* is the total number of phrases, and *P* = *s × t*, for *s* = 306 sensors and *t* time points. Each element of the data matrix, *x*_*i,j*_, represents the value for training instance *i* at a point *j* in MEG sensor/time space. To predict each of the dimensions of the semantic vector, we trained an *L*_2_ regularized (ridge) regression model, 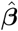:

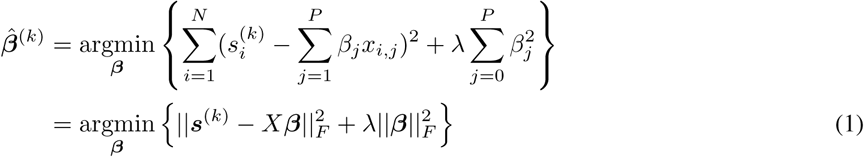

where 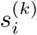 is the *k*th element of the VSM word vector for training instance *i*, and *λ* is a regularization parameter that controls overfitting in ***β***. We append a column of 1s to *X* to incorporate a bias term. Notice that each semantic dimension *k* has its own 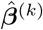. In previous work we have found *λ* to be very stable, so a set value of *λ* = 10^−6^ was used to save computation time. Training data was normalized during cross-validation so that each time-sensor feature has mean 0 and standard deviation 1, and the same correction was applied to test data. Time windows used to train each regression model are 90 ms wide and overlap by 10ms with adjacent windows. Unless specified, all analyses are performed in sensor space.

### 2.5. The 2 vs. 2 Test

For each stimulus word *w* in this study, we have an *m*-dimensional VSM vector 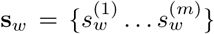, created from corpus data. We refer to each feature 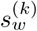 in this vector as the *k*th semantic feature of the stimulus word *w*. The semantic vector may correspond to the adjective or the noun, depending on the analysis being performed.

Using Equation 1, we trained *m* independent functions 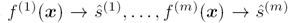, where *ŝ*^(*k*)^ represents the predicted value of the *k*th semantic feature, and 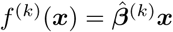 using 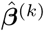 from Equation 1. We combined the output of *f* ^(1)^ … *f* ^(*m*)^ to create the final predicted semantic vector Ŝ = {*ŝ*^(1)^ … *ŝ*^(*m*)^}. We used a distance function to quantify the dissimilarity between two semantic vectors (**Ś** and **s**_**w**_). Many distance metrics could be used, we chose cosine distance due to its popularity for VSM-related tasks in computational linguistics.

To test performance we used the forced choice **2 vs. 2 test** [15]. For each test we withheld the MEG recording for two of the 30 available adjective-noun phrases (phrase *i* and *j*), and trained 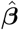 on the remaining 28. We then used the MEG data of the held out phrases (***x***) to predict the semantic vectors for both of the held out phrases (**ŝ**). The task is to choose the correct assignment of predicted vectors **ŝ**_**i**_ and **ŝ**_**j**_ to the true VSM semantic vectors **s**_**i**_ and **s**_**j**_. We will make this choice by comparing the sum of distances for the two assignments:

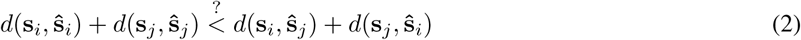

If the left hand side of the above equation is indeed smaller than the left, we mark the 2 vs. 2 test correct. **2 vs. 2 accuracy** is the percentage of correct 2 vs. 2 tests. The 2 vs. 2 test is advantageous because it allows us to use two predictions per test, resulting in higher sensitivity; two weak predictions can still result in a correct assignment of true to predicted phrase vectors. We report the 2 vs. 2 accuracy averaged over all subjects. Under the null hypothesis that MEG data and semantics are unrelated, the expected chance 2 vs. 2 accuracy is 50%.

### 2.6. Correlations between the words of phrases

There is a correlation between the adjectives and nouns in our experimental paradigm. For example, the word “rotten” is always followed by a food word. The word “carrot” never appears with the adjective “gentle”. If we were to not correct for this correlation, we could build a model that was actually detecting the correlated semantics of the noun, when we had intended to build a model that leverages only the semantics of the adjective. To avoid reporting results that rely on this confound, we only consider predicting the adjective or noun when the other word (noun or adjective, respectively) is shared in the 2 vs. 2 pair. That is, when we encounter a 2 vs. 2 pair that contrasts adjectives “rotten” and “big”, we will include it in our analysis only when the *noun is the same for both phrases* (e.g. “rotten tomato” and “big tomato”). Thus, if our prediction framework leveraged the correlated semantics of the noun to predict the adjective, it would be of no use for differentiating between these test phrases. The same is true for the noun; when we encounter a 2 vs. 2 pair that contrasts nouns “dog” and “bear”, we include it only when the adjective is the same for both phrases (e.g. “gentle dog” and “gentle bear”). Thus we can be sure that we are not relying on any correlations between adjectives and nouns for our analyses. There are (30 choose 2) = 435 distinct 2 vs. 2 tests. Amongst those 435 tests, 51 share the same adjective and 60 share the same noun. Our analysis proceeded with these 111 pairs.

### 2.7. Significance Testing

We used permutation tests to determine the probability of obtaining our prediction results by chance. Permutation tests require shuffling the data labels (phrase identity) and running the identical prediction framework (cross validation, training ***β***, predicting *ŝ*, computing 2 vs. 2 accuracy) on the permuted data. When we do this many times, we approximate the null distribution under which the data and labels have no relationship. From this null distribution we calculate a p-value for the performance we observe when training on the true (un-permuted) assignment of words to MEG data. In the experiments that follow, we will train and test multiple predictors on multiple time windows. To account for the multiple comparisons performed over the time windows, we used the Benjamini-Hochberg-Yekutieli (BHY) procedure [25], and *p* = 0.01.

### 2.8. Time Generalization Matrices

To test the consistency of the neural code in time, we use Temporal Generalization Matrices (TGMs) [26]. TGMs mix train and test data from different time windows to test the stability of the neural representation over time. For our TGMs, we used the prediction framework described in Section 2.4, but mix train and test data selected from different time windows for each of the 2 vs. 2 tests. The entry at (*i, j*) of a TGM (*T*_(*i,j*)_) contains the 2 vs. 2 accuracy when we train using MEG data from a time window centered at time *i*, and test using MEG data from a time window centered at time *j*. Thus, depending on the value of *i* and *j* we may be mixing train and test data from different time periods, possibly comparing times when the subject is viewing a different word type, or no visual stimuli at all. If the neural representation of a concept is stable, then models can be trained and tested with data from different time windows with little or no impact on accuracy. TGMs have been use to show that information from previous stimuli can be detected during a memory task, even after stimulus presentation[27, 28]. Our analysis uses windows of the MEG data to compute each *T*_(*i,j*)_, whereas previous work appears to use one time point only. A windowed approach allows us to track the more complex patterns required for language understanding.

## 3.. Results

We will begin with a typical time and ROI analysis of the neural representation of adjective an nouns, and then explore a TGM analysis of our data.

### 3.1. Predicting Adjective and Noun Semantics

2 vs. 2 accuracy for predicting the adjective and noun from whole brain MEG sensor data observed within a sliding 90 millisecond interval appears in Figure 2. Here we can see that the adjective remains detectable until well after the onset of the noun, dipping below statistical significance for the first time at about 1100 ms after the onset of the adjective (300 ms after the onset of the noun). Adjective prediction accuracy again rises above the statistical significance threshold at several subsequent time points, and is significantly above chance for the last time at 1955 ms after the onset of the adjective. Thus, our model can predict the adjective during adjective *and* noun presentation, and also for a prolonged period *after* the noun stimuli has disappeared. This implies that there is a neural representation associated with the adjective that persists during the entire phrase presentation, as well as after the phrase has been presented (we call this the phrase wrap-up period). From these results we cannot tell if the neural representation of the adjective changes over time, we can only infer that there exists a reliable detectable representation during each significantly above-chance time window.

**Figure 2:**
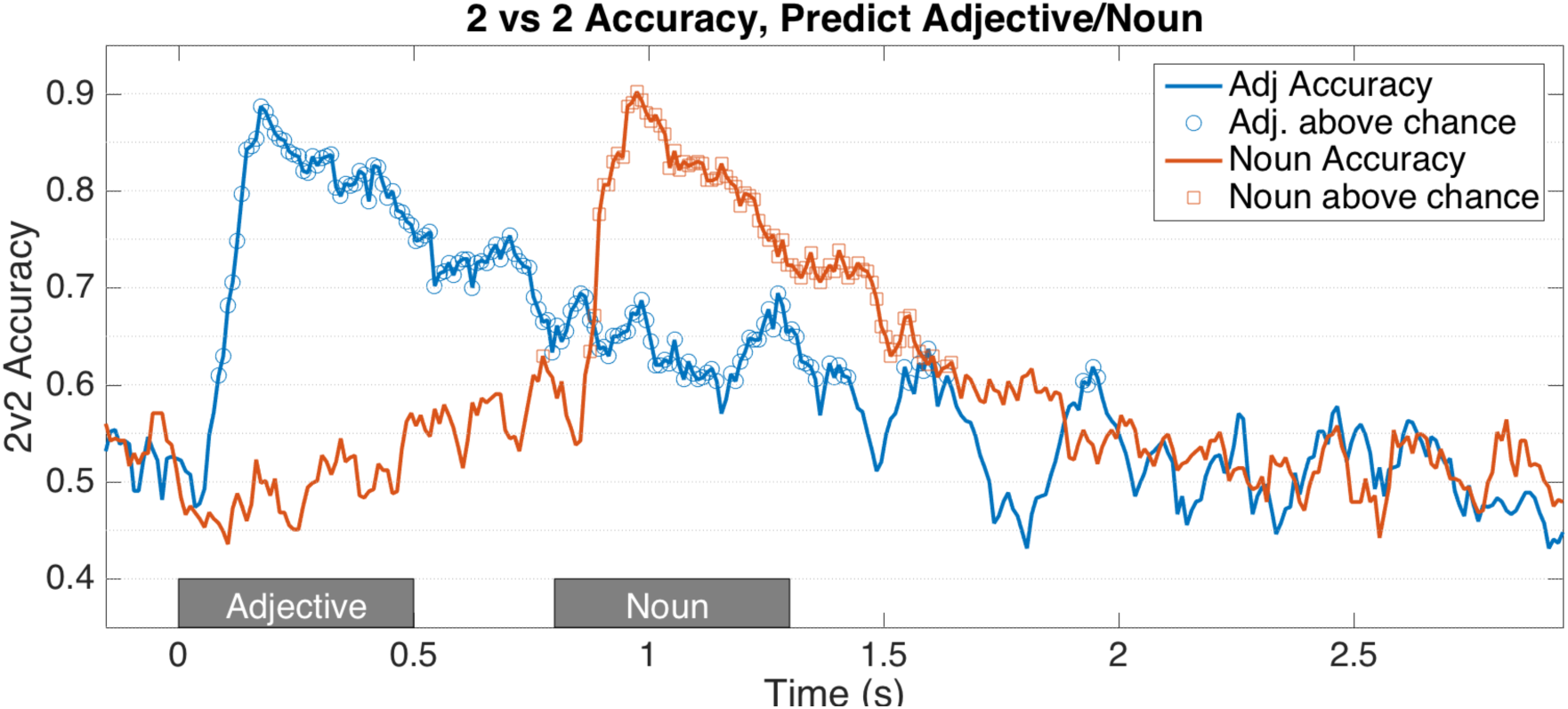
2 vs. 2 accuracy as a function of time for predicting the words of the phrase, averaged over 9 subjects. Time windows for the presentation of the adjective and noun are indicated with grey rectangles. Vertical axis indicates the prediction accuracy for the adjective or noun, based on predicting the word from a 90 millisecond interval of MEG data in sensor space. Significantly above chance accuracies are highlighted. Here a separate prediction model was trained for each time point on the horizontal time axis. Significantly above chance accuracy occurs during word presentation (as expected), but adjective accuracy remains high, even during and after noun presentation.

Figure 2 also shows noun prediction accuracy as a function of time. After its initial peak around 890 ms, the accuracy for predicting noun semantics dips below chance at 1615 ms, and is not above chance after 1645 ms. Thus, the noun is detectable at significantly above chance levels for a continuous interval of size 740 ms. Though prediction accuracy is sustained after the offset of the noun, there is no resurgence later in time. The accuracy for predicting the noun falls off more quickly, and is significant for less time than the adjective. There is also one above chance point very early in time (windows centered at 775 ms, corresponding to time windows 730-820 ms). This window does overlap with the first 20 milliseconds of noun presentation, so may be due to the visual features of the word, which can be correlated to the word vector. Correlation of word vectors to visual features is discussed further in Section 4.3

### 3.2. Localizing Adjective and Noun Semantics

What parts of the brain are driving this late-in-time response to the adjective semantics? To answer this question, we used each subject’s source localized MEG data, parcellated based on the Desikan-Killiany Atlas [29]. All sources localized to a particular ROI were used to train and test a model. In Figure 3, we explore the results for predicting the adjective, and show those ROIs with at least one time point above chance (after FDR correction) in the given time interval. We colored each ROI with the maximum 2 vs 2 accuracy during that time window, obtained with sources localized to that ROI. We explored three time windows: 0.200-0.400 s, 1.545-1.635 s and 1.925-1.955 s. The first time window includes the peak of decoding during the adjective’s presentation, and the last two time windows correspond to the last two time periods the 2 vs. 2 accuracy for predicting the adjective is above chance (Figure 2). We corrected for multiple comparisons over the full 3 s of presentation using BHY FDR correction with p=0.01. A video of 2 vs. 2 accuracy over all ROIs for the full 3 s is available in the Supplementary Material.

**Figure 3:**
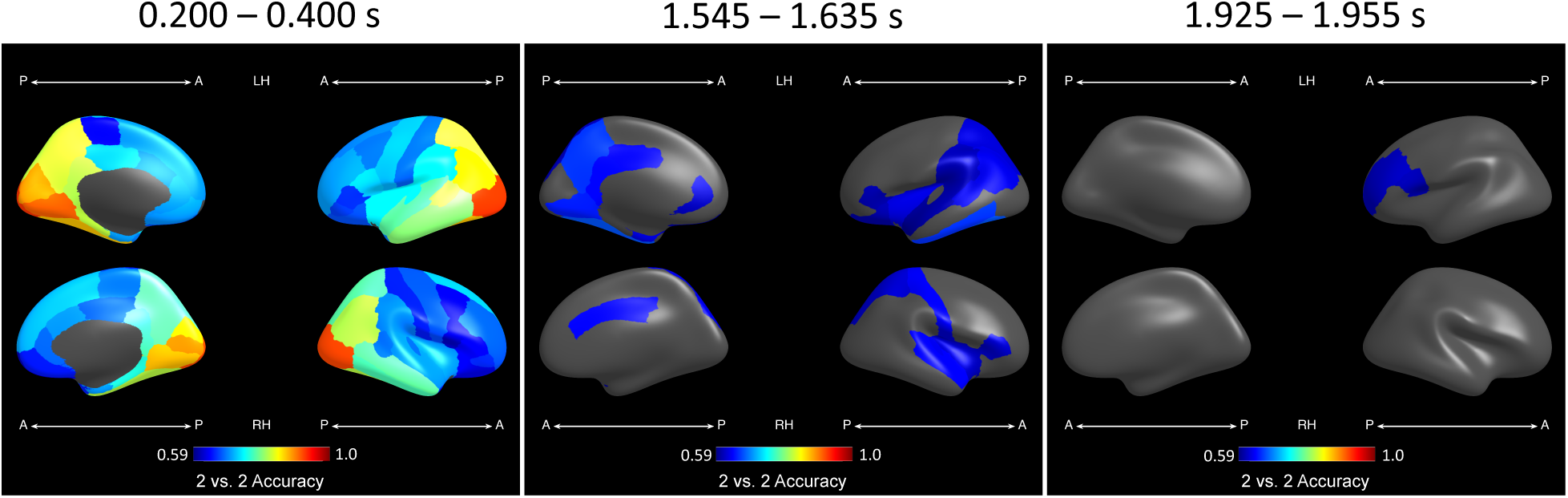
2 vs. 2 accuracy for predicting the adjective using source localized data. The last two time windows correspond to the last two significantly above chance time periods in Figure 2. From left to right, analysis performed over windows centered at: 0.200-0.400 s, 1.545-1.635 s, 1.925-1.955 s. ROIs with no time points above chance during the window are masked, remaining ROIs are colored with the highest 2 vs. 2 accuracy in the corresponding time window. In the earliest time window, during adjective presentation, nearly every ROI can be used to detect the identity of the adjective, with the highest accuracies occurring in occipital and parietal regions. During the later time windows, above chance ROIs appear mostly in parietal, temporal and inferior frontal regions.

As one would expect, across all windows, there are more statistically significant ROIs in the left hemisphere and they have higher peak 2 vs. 2 accuracy. In the earlier time window, 1.545-1.635 s, several temporal and parietal ROIs are above chance, with the highest 2 vs. 2 accuracies occurring in the left fusiform, precuneus, and inferior temporal regions (2 vs. 2 accuracies of 0.67, 0.66 and 0.66 respectively). In the later time window, 1.925-1.955 s, only two regions are above chance: left rostral frontal and left pars opercularis (Broca’s area), both having 2 vs. 2 accuracy 0.6. This late accuracy in Broca’s area lends support to our claim that compositional processes may be occurring as late as 2 seconds after the onset of the adjective, a point we will return to in the discussion section.

### 3.3. Consistency of the Neural Code in Time

How consistent in time is the neural code of the adjective? That is, does the neural code for the adjective during adjective presentation resemble the neural code used for the adjective during noun presentation, or during the phrase wrap-up period? To test this, we created TGMs (Figure 4), as described in Section 2.8, using whole brain MEG sensor data. Recall that each point along the *diagonal* of the TGMs corresponds to training and testing the prediction model using MEG data taken from the same time window, and so represents exactly the results plotted in Figure 2. In contrast, the off-diagonal elements correspond to training the model using MEG data from one time interval, but testing its prediction accuracy on data taken from a second time interval, allowing us to test whether the neural representation of this predicted information varies over time.

**Figure 4:**
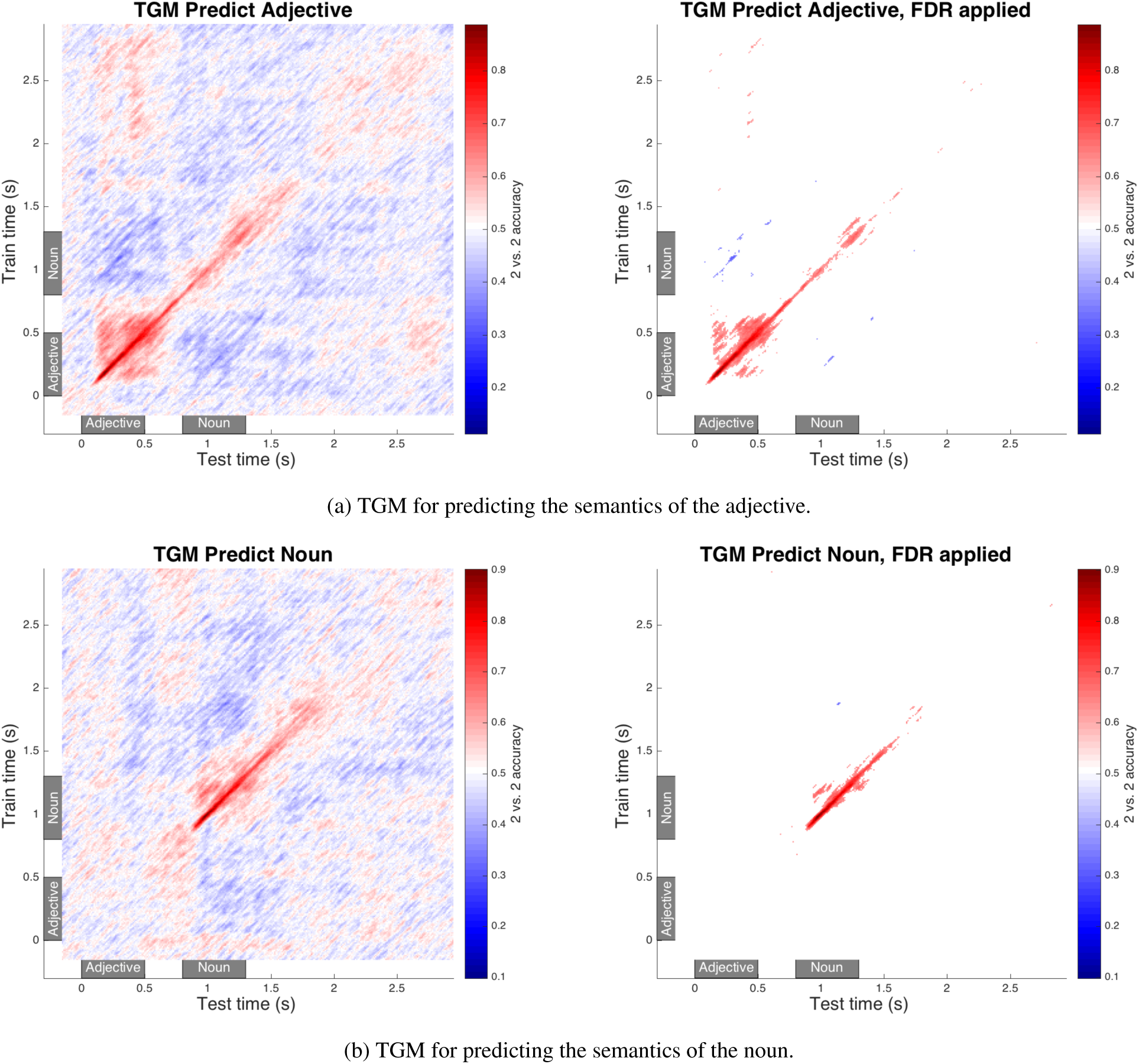
2 vs. 2 accuracy for predicting the words of the phrase, presented in TGMs. Left: all results, Right: FDR thresholded. A TGM mixes training and testing data in all possible combinations to track the similarity of a neural representation in time. Within each TGM, the color at point *i, j* indicates the prediction accuracy when the model is trained using data from an interval centered at time point *i*, then tested by its ability to predict the noun or verb based on MEG data centered at time point *j*. Time windows are 90 ms wide and overlap by 10 ms with adjacent windows. Time 0 is the onset of the adjective, 0.8 is onset of the noun, as annotated with grey rectangles. The adjective shows a resurgence after the presentation of the noun which matches the representation observed during adjective presentation, whereas the noun shows no such resurgence pattern.

The TGMs in Figure 4 show two key features. Firstly, there are off-diagonal patches that correspond to both high and low prediction accuracies. Only the adjective TGM shows off-diagonal accuracy patches above chance (Figure 4a), and they are strongest when training on a time point after 2 s, and testing on a period during the presentation of the adjective. Significantly below chance accuracy patches appear for both the adjective and noun (thought the noun patch is small), and are strongest when training on a window just after the offset of that word and testing during the time when a word is visible (discussed further in Section 4.5). Secondly, both TGMs show a highly oscillatory pattern, which manifests as many diagonal lines parallel to the main diagonal. To quantify this effect, we performed a Fourier transform on the columns of the TGM. We found that TGM oscillations fall into the alpha band family of frequencies (8-13 Hz), but also that they were highly correlated to the amplitude spectrum of the raw MEG data, including the alpha peak.

### 3.4. Results Summary

To summarize, the main findings from our data analyses are: 1) we can predict the adjective well after noun stimuli presentation (as late as 2s after adjective presentation), 2) the neural representation of adjective semantics observed during adjective reading is reactivated after the entire phrase is read, with remarkable consistency, and 3) there is a period of below chance prediction performance just after the presentation of each word.

## 4.. Discussion

Here we compare and contrast the results from Section 3 to build a hypothesis of how the brain represents and processes adjective-noun phrases. To foreshadow, our interpretation of the above analyses for adjective noun processing is as follows:

1. During the time the adjective is read, the brain maintains a neural representation for the adjective.
2. During the time the noun is read, the brain holds both the representation for the noun, and also a representation of the adjective that is the neural “reverse”^1^ of the representation during adjective reading.
3. After the noun stimuli ends, the noun also enters a reversed representation state, though much less pronounced.
4. The adjective’s representation resurges after reading the noun, and it is a good match to the representation observed during adjective reading.
5. Our ROI analysis confirms previous reports of a distributed representation of semantics, and the last resurgence of adjective semantics appears to be localized to left inferior frontal ROIs.

### 4.1. Adjective Semantics in Early and Late Time Windows

Figure 4a (adjective semantics TGM), shows that the pattern for the adjective is fairly consistent within the time that the adjective is being presented. In addition, there are significantly above chance points very late in time, as late as 2.5 s. This result implies that the early and late representations of adjective semantics are highly similar. This late above chance accuracy could be due to the intersective quality of most of the adjectives chosen for this study; the meaning of our selected adjectives is largely unaffected by the semantics of the nouns in this experiment. For example, rotten has very similar meaning when paired with either tomato or carrot. Though the two foods may spoil in slightly different ways, the end result is inedible. Thus, it is reasonable that, for phrases that contain an intersective adjective, the neural representation of the adjective alone should be very similar the phrase’s neural representation. It should be noted, however, that since we did not include a non-compositional condition (e.g. word lists), we cannot definitively say that the results are due to composition.

At first glance, one might think the late above chance prediction accuracy for the adjective conflicts with previous work showing that semantic composition begins as early as 140 ms after the onset of the noun, and as late as 600ms [30, 2]. In our experiments, this would correspond to 940 ms and 1400 ms after the onset of the adjective. The early effects of semantic composition are typically studied using contrasting stimuli that either does or does not require composition. Thus, the timings reported in previous work are the “switching on” of the machinery required to perform semantic composition, but not necessarily the time when we would expect to see the final product of semantic composition, or cessation of thinking about the phrase semantics. Our analysis is specifically looking for the signature of individual words in a composed representation, and thus it is logical that it appears after the typical P600 effect signaling composition violations.

There is support for semantic effects as late as the effects we see here. Previous work has shown effects during joke comprehension as late as 1100 ms after the onset of the final word [31]. Semantic violation effects have been reported as late as 2.5 s after the onset of the critical word in the sentence [6]. When the semantic plausibility of sentence critical words is varied, differences in EEG recordings for anomalous vs expected words extend to the edge of the 1.2s analysis window (and possibly beyond) [32]. Many analyses have restricted themselves to the time period ending 1s after the onset of the critical word, possibly because the paradigms only allowed for analysis to that point [1, 2]. A review of several other accounts of this late activation appears in VanPetten and Luka [33]. The results of the present study show that analyzing MEG data beyond 1s post stimulus onset can give new insight into semantic processing.

One might also wonder if the late resurgence is a task effect. Recall that participants were asked to press a button when they were presented with two adjectives in a row. It is possible to do this task without performing composition, and without attending to the first word at all, as the task can be simplified to pressing the button if the *second* word is an adjective. However, as we analyze only the non-oddball (adjective noun) trials, if a person was only attending to the second word, the late resurgence should show the signature of the noun. Instead, the late resurgence shows the signature of the adjective. Still, the late resurgence could be a byproduct of recalling the first word in order to perform the consecutive adjectives task, rather than a byproduct of adjective-noun composition. This is a disadvantage of our collection paradigm, and future work should incorporate a task that explicitly requires composition.

However, the extremely distributed nature of adjective semantics seen during adjective reading is not replicated in these late time windows (Figure 3). Instead, significant ROIs later in time tend to be confined to areas known to be associated with composition, specifically temporal and left frontal areas. This is particularly striking for the final window, 1.925-1.955 s, where the only areas above chance are in the left frontal lobe, and include Broca’s area. This is in line with Hagoort’s theory of composition [3], which place unification areas in LIFG, and control areas in dorsolateral prefrontal cortex.

### 4.2. Noun Semantics

The neural representation of the noun is detectable until 1645 ms after the onset of the adjective (845 ms after the onset of the noun). This duration is shorter than that of the adjective, which is detectable in the brain as late as 2 s after the onset of the adjective stimulus. After the noun stimulus has left the screen, a “reversed” representation of the noun appears in the brain, again for a much shorter time than the adjective.

Noun semantics are not predictable during the phrase wrap up period (1.3-3s). It is somewhat counter-intuitive that the semantics of the adjective should be more salient than the semantics of the noun during the contemplation of the phrase. This could be the result of our choice of adjectives, which manipulate the most predictable features of the noun to their extreme ends, perhaps obfuscating the prototypical noun representation.

### 4.3. Decoding Visual Features

Some critiques of a decoding approach for studying semantics claim we may only be detecting the visual features of the word form, not the semantics. To address this concern, we ran our analysis to predict the number of letters in the adjective, a feature highly correlated with the visual information available to the participant. In a sensor space analysis, we found that after 700ms, we could not reliably predict word length. Word length was most predictable in occipital cortex ROIs before 200ms post stimulus onset, and not predictable later in time. Note also that in Figure 4a, high off-diagonal accuracy appears for *test* times at 200ms and later, so is likely not attributable to a visual feature of the stimulus being recalled. In addition, TGMs for word length contained no significant off-diagonal points, implying that the resurgence of adjective semantics is not simply the image of the word being visually recalled.

Recall, also, that we are using a corpus-derived semantic representation of the adjective for the prediction tasks throughout this paper. Though there are some correlates to the perceptual features of word strings in these corpus-derived features (e.g. frequent words are, on average, shorter than infrequent words) we are, by and large, leveraging the semantic features of the words when we use these vectors.

### 4.4. The Localization of Semantic Representations

What brain regions are driving the late adjective decodability? The 2 vs. 2 results in the 1545-1635 ms window are largely driven by temporal and parietal regions. This implies that the output of semantic composition involves both distributed parietal regions (as seen for single words in [16]) as well as a contribution from ROIs involved in semantic composition of adjective noun phrases (LIFG, LATL) [4, 5], and specifically intersective adjective noun phrases (LATL) [34]. The later time window (1925-1955 ms) shows above chance decoding in rostral and pars opercularis (Broca’s Area). Broca’s area has long been associated with semantic composition, and is important in several models of semantic composition [35, 3].

### 4.5. Significantly Below Chance Prediction Accuracy in Temporal Generalization Matrices

For both words of the phrase, the TGMs shows a period of below chance prediction accuracy after the offset of the word stimuli. Significantly below chance accuracy may seem counter-intuitive; how can the framework’s predictions be systematically *worse* than random guessing? If the prediction is systematically inverted, perhaps the MEG signal itself is reversed or negated. To test this, we negated the original MEG signal (multiplied by −1) in TGM coordinates that are below chance. We found that the 2 vs. 2 accuracy of negated data was not only *above* chance, it was exactly 1 *− a*, where *a* is the 2 vs. 2 accuracy on the original MEG data. This is a byproduct of our prediction framework. Negated MEG signal leads to negated predictions (that is a negated 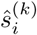 from Equation 1), which causes the predicted vector to point in exactly opposite of the prediction for non-negated data. The opposing direction results in a negation of the cosine of the angle, and thus flips the decision in Equation 2. This negated representation of the word could be how the brain to maintains context while processing a new word.

What does it mean for the MEG signal to be reversed or negated? MEG measures the post-synaptic potential of many parallel dendrites. Thus, the negation of the MEG signal equates to the reversal of the underlying magnetic field. In pyramidal neurons, this could indicate that the opposite end of the same neuron is receiving input, indicating that the same brain area is receiving input, but the source of the input has changed.. This could be caused by several phenomena, perhaps related to the neural loops currently thought to be related to neural oscillations [36]. It is interesting that the negated representation for the adjective appears during the time we would expect to see N400 and P600 effects during noun reading (approximately 1.2 and 1.4 s in Figure 4a).

We hypothesize that for both the adjective and the noun, the brief “reversed” representation is a holding pattern that allows the brain to store a word’s meaning in memory while performing another action. In the case of the adjective, the next action is reading the noun. In the case of the noun, the next action is recalling the adjective for composition. The beginning of the negated noun representation aligns well with the end of the negated adjective representation. After the negated noun representation completes, we see the return of the adjective in its original, non-negated, form. Further studies will be needed to confirm or deny this hypothesis.

### 4.6. The Oscillatory Nature of TGMs

One of the most striking patterns in the TGMs is the oscillatory nature of the prediction accuracy (Figure 4). The oscillation frequency and intensity is different for each participant (TGMs for individual subjects appear in Appendix, Figures A1 and A2), but always resides in the alpha range. Initially we thought this might be a clue that semantic representations repeat themselves, but we noted that the frequencies of the oscillations in the TGMs were highly correlated to the main frequencies in the raw MEG data. Thus, it is possible that the oscillations may simply be an artifact of alpha waves superimposed on the signal corresponding to the semantic representation. In addition, our unjittered stimuli may have exasperated this effect, as alpha entrains to visual signals []. This is a cautionary tale for the subtleties of interpreting TGMs. Because the analysis crosses time boundaries, drifts and oscillations in the signal can have a large effect on results, even when they are not apparent in typical analyses in which the train and test windows are the same.

Still, the correlation of TGM oscillation frequency to subject-specific alpha does not rule out the possibility that semantic representations are oscillatory. Certainly there is support for oscillations playing a role in scene understanding [36], and evidence for alpha-entrained neural signals for visual tasks (perceptual echo) [37]. Further work is needed to explore the role of oscillatory activity in semantic composition.

## 5.. Conclusion

This paper conveyed several new findings regarding adjective-noun composition in the human brain as we tracked the flow of information during semantic composition. Our analysis showed that adjective semantics are predictable for an extended period of time, almost continuously until 1.6s after the onset of the adjective, and are reactivated during late processing, 2-3s after the onset of the adjective (1.2-2.2s after the onset of the noun). The reactivated neural representation matches the representation seen during the initial reading of the adjective. After the offset of each word, a “reversed” representation of the word appears.

The resurgence of adjective semantics is much later than the activation of the machinery responsible for combinatorics, as documented in previous research [4, 7]. The combinatorial machinery of LATL and LIFG could be the hub that coordinates areas, readying them for the compositional processing. This would require them to activate sooner than areas of the brain that *store* the composed semantic meaning. Our results imply that future research interested in the composed representation should look beyond the typical 1s time window after the onset of a word.

With respect to semantic composition in the brain, several new research questions have emerged. For example, does adjective resurgence appear even for non-intersective adjectives? We would also like to explore more complex composition tasks like sentences, paragraphs, stories, and beyond. We are interested in the underlying mechanisms that give rise to significantly below chance accuracy, and would like to explore their role in compositional processing. Our work also raises new questions regarding the role of oscillations in the neural processing of language. By exploring simple composition in a controlled setting, our study establishes an analysis framework for such future research directions.

## Acknowledgements

This material is based upon work supported by the National Institutes of Health grant number 1r01hd075328-0, IARPA afrl award #31081.1 and the CNBC Multimodal Neuroimaging Training Program Fellowship. Alona Fyshe is supported by CIFAR (Canadian Institute for Advanced Research) and NSERC (Natural Sciences and Engineering Research Council).

## Stimuli

The phrases for the adjective noun brain imaging experiment are made from 6 nouns (“dog”, “bear”, “tomato”, “carrot”, “hammer”, “shovel”) and 8 adjectives (“big”, “small”, “ferocious”, “gentle”, “light”, “heavy”, “rotten”, ‘‘tasty”), as well as two null words: “the” and “thing”. The phrases are:

- the dog
- the bear
- the tomato
- the carrot
- the hammer
- the shovel
- big dog
- big bear
- big tomato
- big carrot
- big hammer
- big shovel
- small dog
- small bear
- small tomato
- small carrot
- small hammer
- small shovel
- ferocious dog
- ferocious bear
- gentle dog
- gentle bear
- light hammer
- light shovel
- heavy hammer
- heavy shovel
- rotten carrot
- rotten tomato
- tasty carrot
- tasty tomato

Due to multiple word senses, the word “light” was not used in the adjective-adjective oddballs.

## Word Vector Model

The word vector model used here is based on previous work [24]. It was built using a large corpus of text from web pages, specifically a 16 billion word subset of ClueWeb09 [38]. The sentences were then dependency parsed and statistics were calculated to model the probability of seeing two words in a particular dependency relationship in a sentence. These probabilities were arranged in a large matrix, which was then compressed using Singular Value Decomposition. The first 100 dimensions of the matrix were used for this study.

## TGM analysis for all subjects

**Figure A1:**
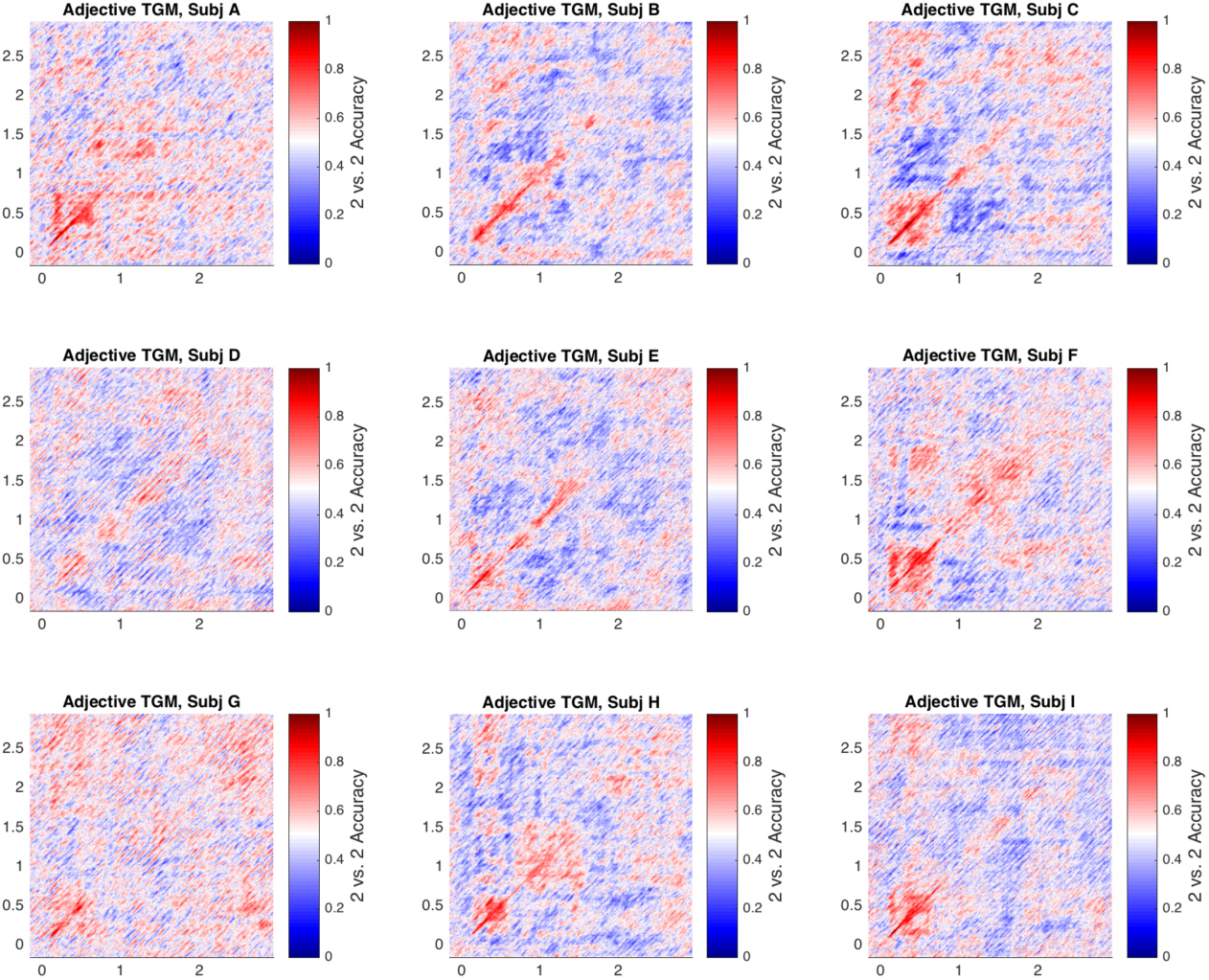
The Adjective TGM matrices for all 9 subjects included in the study.

**Figure A2:**
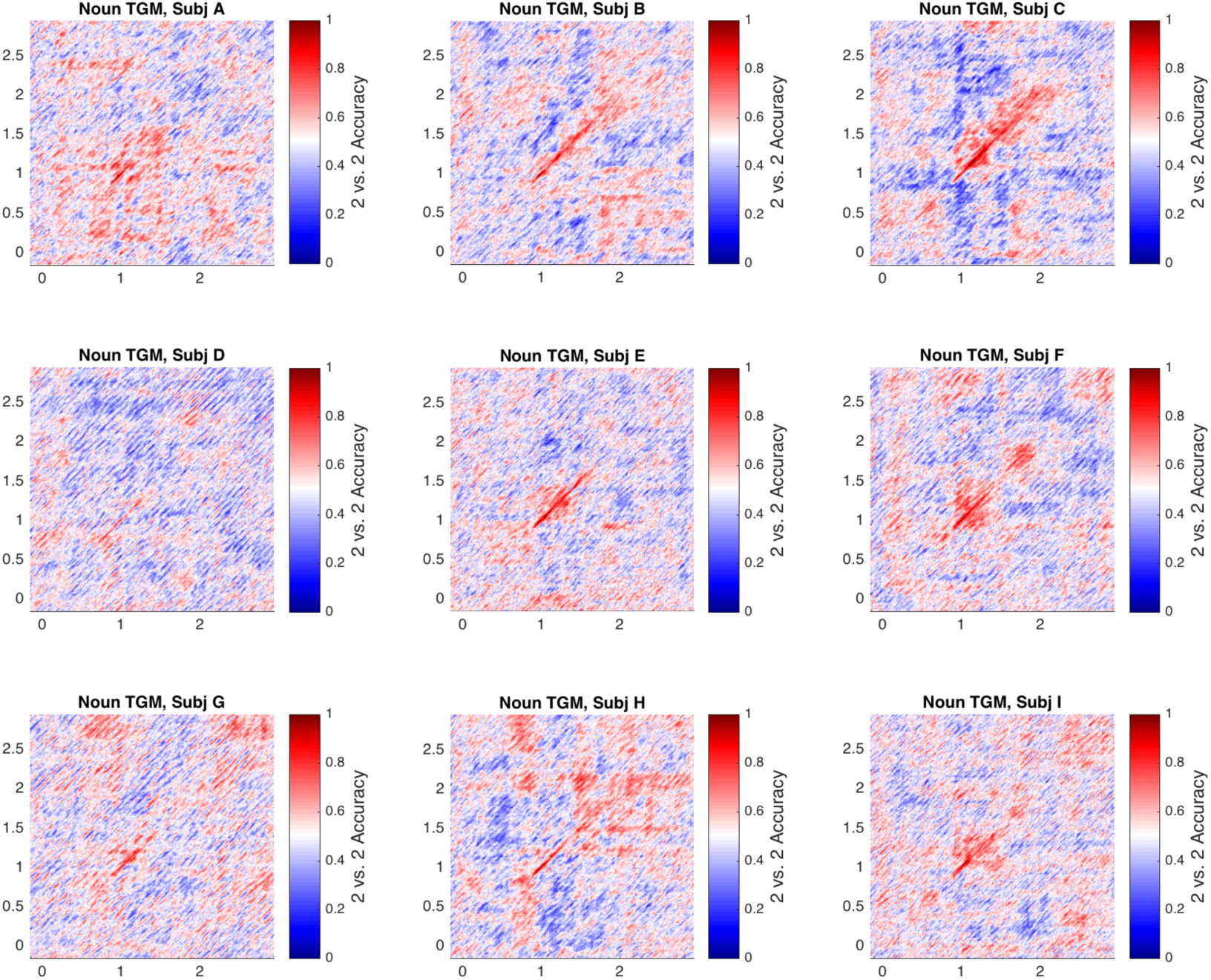
The Noun TGM matrices for all 9 subjects included in the study.

## Footnotes

1 This terminology is borrowed from [26]

